# Brain functional connectivity modulates social bonding in monogamous voles

**DOI:** 10.1101/752345

**Authors:** M. Fernanda López-Gutiérrez, Zeus Gracia-Tabuenca, Juan J. Ortiz, Francisco J. Camacho, Larry J. Young, Raúl G. Paredes, Nestor F. Diaz, Wendy Portillo, Sarael Alcauter

**Affiliations:** Instituto de Neurobiología, Universidad Nacional Autónoma de México, Querétaro, México; Silvio O Conte Center for Oxytocin and Social Cognition, Center for Translational Social Neuroscience, Yerkes National Primate Research Center, Department of Psychiatry and Behavioral Sciences, Emory University, Atlanta, Georgia 30329, United States of America; Instituto Nacional de Perinatología Isidro Espinosa de los Reyes, Mexico City, México

## Abstract

Previous studies have related pair bonding in *Microtus ochrogaster*, the prairie vole, with plastic changes in several brain regions. However, their socially-relevant interactions have yet to be described. In this study, we used resting state magnetic resonance imaging to explore longitudinal changes in functional connectivity of brain regions associated with pair bonding. Male and female prairie voles were scanned at baseline, after 24 hours and two weeks of cohabitation with mating. Network based statistics revealed a common network with significant longitudinal changes including prefrontal and cortical regions, the hippocampus, the anterior olfactory nucleus, the lateral septum, the paraventricular nucleus, and the ventral tegmental area.

Furthermore, baseline functional connectivity of three sub-networks predicted the onset of affiliative behavior, and a relationship was found between partner preference with long-term changes in the functional connectivity between the medial amygdala and ventral pallidum. Overall, our findings revealed the association between network-level changes and social bonding.

## Introduction

The prairie vole (*Microtus ochrogaster*) is a rodent native of North America whose natural behavior involves pair bonding, which can be defined as a long-lasting, strong social relationship between individuals in a breeding pair in monogamous species (Walum & Young, 2018). Pair bonded voles will usually display selective aggression towards unfamiliar conspecifics; biparental care, including paternal behavior and alloparenting; incest avoidance and reproductive suppression of adult individuals within a family group (Carter et al., 1995). These behaviors make the prairie vole a valuable model to investigate behaviors associated with a socially monogamous reproductive strategy (L. J. Young & Wang, 2004), social bond disruption, social isolation and social buffering (Lieberwirth & Wang, 2016). Being comparable to human-like social interactions, studying the neurobiology of social behavior in the prairie vole may allow further understanding of human social bonding, its alterations in psychological disorders and overall impact in health (Kiecolt-Glaser et al., 2010).

Pair bonding-related behaviors in the prairie vole depend on hormonal mechanisms and the activation of emotional, reward and sensory brain circuits (Walum & Young, 2018), integrating large functional networks (Johnson & Young, 2017). Among these, the mesolimbic reward system and the social decision-making network (SDMN) (Johnson et al., 2017) are the proposed networks to be involved in pair bonding, being modulated by steroid hormones, dopamine (DA), oxytocin (OXT), arginine vasopressin (AVP), γ-aminobutyric acid (GABA), glutamate, and corticotropin-releasing factor (CRF), among others (Walum & Young, 2018). A current hypothesis suggests that pair bonding consists of two different plastic processes: the formation of a neural representation of the partner, and a selective preference for the partner, i.e. the maintenance of the pair bond (L. J. Young & Wang, 2004; Walum & Young, 2018). In this process, an association has to be created between the reinforcing properties of sex (mating) and the olfactory signature from the partner (Ulloa et al., 2018; L. J. Young & Wang, 2004). In broad terms, both OXT and AVP are necessary and sufficient for the formation of the pair bond (L. J. Young & Wang, 2004; Lieberwirth & Wang, 2016), and their release during social and sexual interactions are the likely triggers and critical modulators of the mentioned network, since most regions included in the SDMN express their corresponding receptor binding (Johnson & Young, 2017). DA would also be released in concert with OXT and AVP in specific regions to modulate the adequate display of behavior and formation of the pair bond (L. J. Young & Wang, 2004), including the NAcc, which has substantially more D1-like receptor binding two weeks after female exposure, relative to non–pair bonded males (Aragona et al., 2006). However, the interplay of these broad networks has not been directly explored *in vivo*.

Recently, novel electrophysiologic and optogenetic techniques have been employed to demonstrate that the functional connectivity between the nucleus accumbens (NAcc) and the medial prefrontal cortex (mPFC) during initial cohabitation in female prairie voles modulates the affiliative behavior with their potential partner (Amadei et al., 2017), providing exciting data of the relevance of such corticostriatal interactions for social bonding. However, this approach does not allow the study of the interaction of multiple brain regions, i.e. networks, and their relevance of such interactions in the process of pair bonding. Neuroimaging methods may provide the alternative to explore such networks in a longitudinal fashion, since few studies have made use of positron-emission tomography (PET) to explore limited aspects of such longitudinal changes (Bales et al., 2007), providing the first longitudinal evidence of neurophysiological changes associated to pair bonding. Potentially, functional magnetic resonance imaging (fMRI) may be the ideal tool to explore the longitudinal changes in the functional brain networks (Damoiseaux et al., 2006), providing high spatial resolution, wide brain coverage and being minimally invasive to explore longitudinal changes. In particular, resting state functional magnetic resonance imaging (rsfMRI) explores the low frequency fluctuations (<0.1□Hz) of the blood oxygen level dependent (BOLD) signal, which has proven to remain highly synchronized within the sensory, motor and associative networks in the brain of humans (Damoiseaux et al., 2006), non-human primates (Rilling et al., 2007) and rodents (Gozzi & Schwarz, 2016), including the prairie vole (Ortiz et al., 2018), turning this technique into a promising tool for translational research. Indeed, functional connectivity explored by means of the correlation of rsfMRI signals of anatomically separated brain regions (Friston et al., 1993) has demonstrated to be correlated with neuronal activity (Mateo et al., 2017), and it is subject to change in plastic processes such as learning and memory in both humans (Jolles et al., 2013) and rodents (Nasrallah et al., 2016). Here, we make use of this non-invasive technique to explore the longitudinal changes in the brain functional connectivity associated with pair bonding in prairie voles.

## Materials and Methods

### Animals

Thirty-two three-month-old sexually naïve female (*N*=16) and male (*N*=16) prairie voles (*Microtus ochrogaster*) were used in the study. The animals were housed in a temperature (23°C) and light (14:10 light-dark cycle) controlled room and provided with rabbit diet HF-5326 (LabDiet, St. Louis, MO, USA) oat, sunflower seeds, and water *ad libitum*. These voles were previously weaned at 21 days, housed in same-sex cages, and were descendants of voles generously donated by Dr. Larry J. Young from his colony at Emory University. The number of subjects per group is comparable to the largest sample sizes using rsfMRI in rodents (Bajic et al., 2016; Christiaen et al., 2019; Grandjean et al., 2014). All surgical, experimental and maintenance procedures were carried out in accordance with the “Reglamento de la Ley General de Salud en Materia de Investigación para la Salud” (Health General Law on Health Research Regulation) of the Mexican Health Ministry which follows the National Institutes of Health’s “Guide for the Care and Use of Laboratory Animals” (NIH Publications No. 8023, revised 1978). The animal research protocols were approved by the bioethics committee of the Instituto de Neurobiología, UNAM.

### Surgical procedures

Fourteen days before the experimental protocol, female voles were bilaterally ovariectomized. After recovery, silastic capsules (Dow Corning™ Silastic™ Laboratory Tubing; Thermo Fisher Scientific, Pittsburg, USA) containing estradiol benzoate (E2B; Sigma Aldrich, Missouri, USA) dissolved in corn oil (0.5 mg/mL of E2B) were implanted via s.c. to induce sexual receptivity four days before cohabitation protocol and remained implanted during the entire experimental protocol. The corresponding dose and procedure reliably induced sexual receptivity (Ingberg et al., 2012).

### Anesthesia for image acquisition

Animals were anesthetized to avoid stress and excessive movement during scanning sessions. Isoflurane at 3% concentration in an oxygen mixture was used for induction and positioning in the scanner bed, in which the head was immobilized with a bite bar and the coil head holder. Once voles were securely placed in the scanner bed, isoflurane anesthesia was adjusted at a 2% concentration and a single bolus of 0.05 mg/kg of dexmedetomidine (Dexdomitor; Zoetis, Mexico) was administered subcutaneously. Five minutes after the bolus injection, isoflurane anesthesia was lowered and maintained at 0.5%. MRI acquisition started when physiological readings were stabilized (~15 minutes after bolus injection). The use of both anesthetics has been reported as optimal for rsfMRI acquisition in rodents and closely resemble an awake condition (Grandjean et al., 2014, Paasonen et al., 2018), yet the mentioned combination of dose and administration route were previously standardized for this specific protocol in prairie voles (non-published data). Body temperature was maintained with a circulating water heating pad within the scanner bed, respiration rate was monitored with an MR-compatible pneumatic pillow sensor, and blood oxygen saturation was measured with an MR-compatible infrared pulse-oximeter (SA instruments Inc, Stony Brook NY). After the scanning sessions, animals were monitored until fully recovered and transferred back to their housing.

### Image acquisition

Prairie voles underwent three MRI acquisition sessions: a baseline scan before cohabitation, a second scan after 24 hours of cohabitation, and a third scan after two weeks of cohabitation. The timeline of the experiment is described in Figure 1. MRI acquisition was conducted with a Bruker Pharmascan 70/16US, 7 Tesla magnetic resonance scanner (Bruker, Ettlingen, Germany), using an MRI CryoProbe transmit/receive surface coil (Bruker, Ettlingen, Germany). Paravision-6 (Bruker, Ettlingen, Germany) was used to perform all imaging protocols. Before running the fMRI sequence, local field homogeneity was optimized within an ellipsoid covering the whole brain and skull using previously acquired field maps. rsfMRI was acquired using a spin-echo echo planar imaging (SE-EPI) sequence: repetition time (TR) = 2000 ms, echo time (TE) = 19 ms, flip angle (FA) = 90°, field of view (FOV) = 18 × 16 mm^2^, matrix dimensions = 108 × 96, yielding an in-plane voxel dimensions of 0.167 × 0.167 mm^2^, and slice thickness of 0.7 mm, total volumes acquired = 305 (10 minutes and 10 seconds). After the rsfMRI sequence, an anatomical scan was obtained using a spin-echo rapid acquisition with refocused echoes (Turbo-RARE) sequence with the following parameters: TR = 1800 ms, TE = 38 ms, RARE factor = 16, number of averages (NA) = 2, FOV = 18 × 20 mm^2^, matrix dimensions = 144 × 160, slice thickness = 0.125 mm, resulting in isometric voxels of size 0.125 × 0.125 × 0.125 mm^3^.

**Figure 1.**
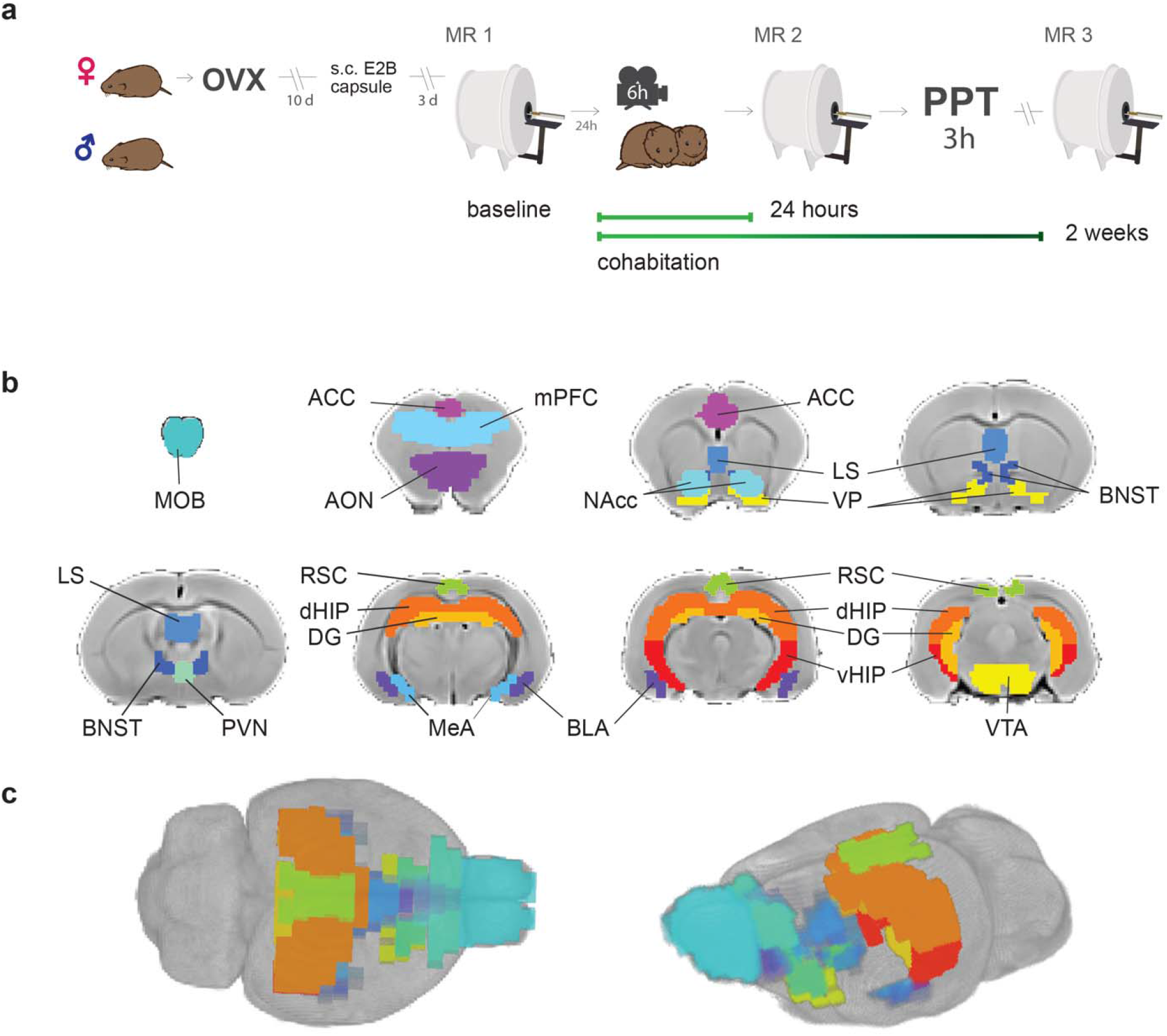
a) Sequence of experiments during a 30-day period: Female voles were bilaterally ovariectomized before MR and behavioral protocols. After being allowed to recover from surgery for 10 days, silastic capsules containing E2B (estradiol benzoate) were implanted via s.c. 4 days for sexual receptivity induction before cohabitation. Once couples went under cohabitation, they were housed together for the remaining of the experiment and were only separated for PPT and MR scanning sessions. OVX: ovariectomy surgery. MR: magnetic resonance imaging scanning session. PPT: Partner preference test. **b)** Regions of interest (ROIs) for network functional connectivity analysis. Antero-posterior coronal slices of the prairie vole template overlayed with ROI masks with the resolution used in the analysis. Each color represents a different ROI. ACC: anterior cingulate cortex. AON: anterior olfactory nucleus. BLA: basolateral amygdala. BNST: bed nucleus of the stria terminalis. DG: dentate gyrus. dHIP: dorsal hippocampus. MeA: medial amygdala. MOB: main olfactory bulb. LS: lateral septum. mPFC: medial prefrontal cortex. NAcc: nucleus *accumbens*. PVN: paraventricular nucleus. RSC: retrosplenial cortex. VP: ventral pallidum. vHIP: ventral hippocampus. VTA: ventral tegmental area. **c)** 3D views of ROI masks embedded within the prairie vole template.

### Cohabitation and behavior analysis

The day after the baseline scanning session, female and male voles unrelated to each other were randomly assigned as couples and placed for cohabitation in a new home cage with fresh bedding to promote *ad libitum* mating and social interaction. The first 6 hours of cohabitation were video recorded for subsequent analysis of social and mating behavior. Specifically, mount, intromission and ejaculation latencies from male voles; lordosis latency from females; and huddling latencies for each male and female pair. Huddling behavior was considered when the pair was sitting or lying in bodily contact for at least 10 seconds. Voles were housed in couples for the remaining of the experiment and were only separated for MRI scanning sessions and behavioral tests.

### Partner Preference Test

Each subject underwent a 3-hour partner preference test to evaluate pair bond formation. This protocol was based on a previously described test (Williams et al., 1992) and was performed in custom built, 3-chambered clear plastic arenas divided by perforated clear plastic barriers that allowed visual, auditory and olfactory contact, but not physical interaction or mating behavior between subjects. In the central chamber, the vole being tested could roam freely, and time spent in the incentive areas at opposite sides of the chamber was recorded. The incentive areas were defined as the proximal space next to the chambers with its “partner” or with an opposite-sex, novel “stranger” vole. All stranger voles were unrelated to subjects in the test and had the same age and hormonal condition than the sexual partner. On each test, partner or stranger voles were randomly and alternately positioned on the opposite chambers of the arena. Male and female partners were tested in alternate time periods (48 or 72h), assigned at random, to enable rest between tests and avoid excessive stress. PPT data analysis was performed with *UMATracker* software (Yamanaka & Takeuchi, 2018), which allowed quantification of the proportion of time spent with each of the stimulus voles. A partner preference index was calculated for each subject, consisting on the time with partner divided by the total time with any of the stimulus voles (time with partner plus time with stranger vole).

### Imaging data preprocessing

Imaging data preprocessing was performed with FMRIB’s Software Libraries (FSL, v5.0.9; Jenkinson et al., 2012). To avoid initial signal instability, the first 5 volumes of each functional series were discarded. The, slice-timing correction and motion correction were applied, using the first non-discarded volume as reference. The reference volume was also taken to determine the rigid-body transformation to the corresponding anatomical image. The resulting rigid-body transformation was combined with a non-linear transformation to a prairie vole brain template obtained from previous published work (Ortiz et al., 2018). Functional images were later warped to the brain template and resampled to a resolution of 0.4 × 0.4 × 0.4 mm^3^. To minimize physiological confounds, the first 5 eigen-vectors (time-series) within the combined non-grey matter mask were obtained (Behzadi et al., 2007). These eigenvectors and the 6 motion parameters (3 rotations, 3 displacements) were regressed out from each subject’s functional series. Datasets were band-pass filtered to retain frequencies between 0.01 and 0.1 Hz (Gorges et al., 2017). Finally, smoothing was applied with a gaussian kernel with a full width at half maximum of 0.8 mm.

### Functional connectivity analysis

To explore changes in functional connectivity in regions associated with socio-sexual behavior before and after cohabitation, sixteen brain regions of interest (ROI) were defined according to their previously reported relevance in the process of pair bond formation and maintenance (Johnson & Young, 2017; Lieberwirth & Wang, 2016; Walum & Young, 2018) (Figure 2). Specifically, ROIs were manually defined on the anatomical prairie vole brain template, visually guided with the Allen Mouse Brain Atlas (Lein et al., 2007), to select the minimum possible number of voxels that included/covered each of the following regions: the anterior cingulate cortex (ACC), anterior olfactory nucleus (AON), basolateral amygdala (BLA), bed nucleus of the stria terminalis (BNST), lateral septum (LS), medial amygdala (MeA), main olfactory bulb (MOB), medial prefrontal cortex (mPFC), nucleus accumbens (NAcc), retrosplenial cortex (RSC), paraventricular nucleus of the hypothalamus (PVN), ventral pallidum (VP), ventral tegmental area (VTA), dentate gyrus (DG), dorsal hippocampus (dHIP) and ventral hippocampus (vHIP). For each subject and session, connectivity matrices were calculated as follows: first, the average time series for each region were extracted from the preprocessed fMRI datasets and then, partial correlation estimates were obtained for all possible pairs of ROIs, partialling out the remaining time series. Finally, correlation values were Fisher’s z-transformed and further tested to identify connected nodes (ROIs) with significant longitudinal changes. Due to technical problems, two subjects missed the baseline MRI acquisition (session 1), and two subjects missed the 2 week-cohabitation MRI acquisition (session 3). Linear mixed models (LMM) were fitted considering sex (male, female) as a fixed variable, and session (baseline, 24 hours and two weeks of cohabitation) and sex-session interaction as random variables. Sets of connected nodes, i.e. networks, with significant sex, session or interaction effects were identified based on the Network Based Statistics (NBS) analysis (Zalesky et al., 2010), implemented in the “Network Based Statistics in R for Mixed Effects Models” package (NBR, https://CRAN.R-project.org/package=NBR), estimating the statistical significance of the identified networks by comparing it with a null distribution obtained with 5000 permutations of the original data (naturally controlling for the size of the original matrices in the permutation tests). In order to identify potential relationships between behavior and functional connectivity, huddling latencies and partner preference indexes were tested as linear covariates of the functional connectivity data in subsequent NBS analyses.

**Figure 2.**
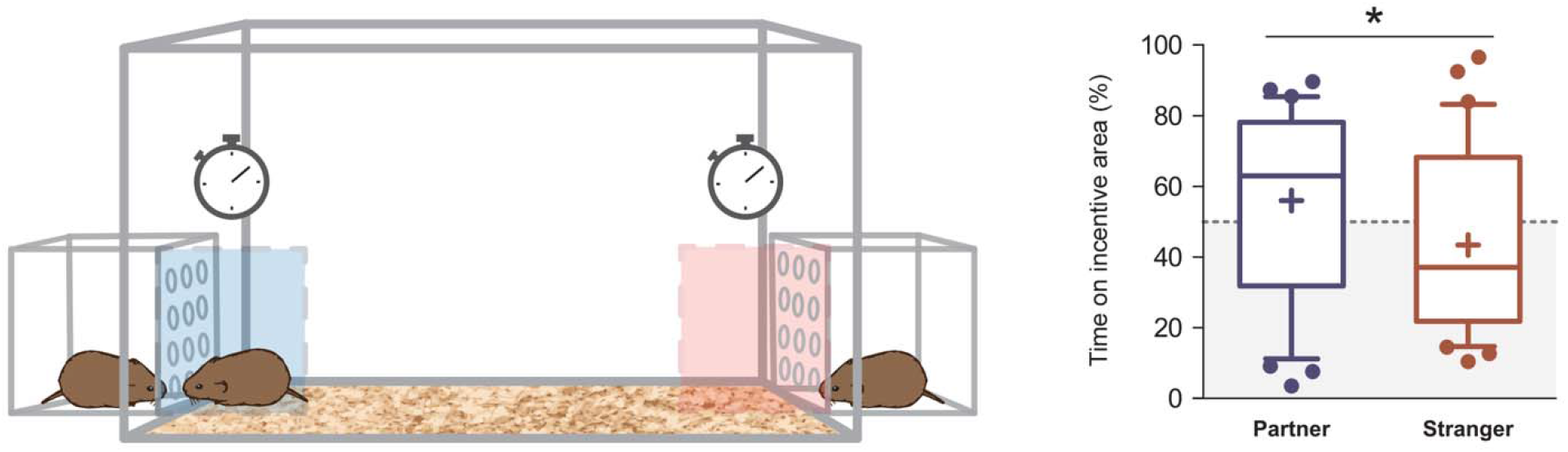
PPT analysis: Between 48 and 72h of cohabitation, partner preference was evaluated on each subject (*N* = 32). A significant difference was found between the time spent on the incentive area related to the partner, with the incentive area related to the stranger vole. Boxplot graphs show whiskers with 10-90 percentiles, horizontal line inside the box shows data median and “+” represents data mean. (*) denotes significance at p < 0.05.

### Statistical analysis

Data in the study is presented as *mean* ± *standard error of the mean* unless otherwise noted. NBR is an R package (https://CRAN.R-project.org/package=NBR) based on Network-Based Statistics, developed by Zalesky and cols. (2010), aimed to identify networks that comply with a statistical test and which size is statistically significant when compared with the null distribution created by permuting the original data. The advantage of NBR is that it allows the implementation of mixed effects models, specially useful for non-balanced longitudinal datasets as the one here analyzed. LMM was computed with ‘nlme’ R package (Pinheiro et al., 2017) to identify significant effects of session, sex or their interaction and integrated with the permutation tests. Post-hoc tests were performed to better describe the significant differences between sessions for each edge in the identified subnetworks, corrected for a false discovery rate (FDR) of 0.05. NBR was also used to find networks with significant linear relationships with behavioral data and *posteriori* Pearson two-tailed correlation tests, obtained with R software package ggplot 2 (ggplot2, RRID:SCR_014601), were used to better describe such relationships for each edge of the subnetworks.

Partner preference was explored with one-tailed Mann Whitney U tests given that Shapiro-Wilk normality tests revealed such data was not normally distributed. Behavioral data was analyzed with GraphPad Prism 5 (*GraphPad* Software, La Jolla, California, USA).

## Results

Mount (*M±SEM:* 65.4±31.7 min), intromission (116±35.3 min), and ejaculation (125±34.4 min) latencies were obtained for male voles (*N*=16). Lordosis latency (22.3±13.3 min) was also measured on females (*N*=16), and huddling latencies (69.5±15.8 min) were obtained for each male and female pair. Three of the sixteen couples did not mate during the recorded period, but all voles displayed huddling and licking/grooming behavior with their sexual partner.

Between 48 and 72 hours of cohabitation, partner preference was evaluated on each subject (*N*= 32) (see methods). A significant difference was found between the percentage of time spent on the incentive area related to the partner (*median* = 55.9 %) with the area related to the stranger vole (*median* = 37.1 %) for all subjects (*U* = 378, *p* = 0.036, *effect size r* = 0.32) (Figure 2). No significant differences were found between males and females in their preference for the partner (*U* = 121, *p* = 0.81, effect size *r* = 0.05) or the stranger voles (*U* = 118, *p* = 0.72, *effect size r* = 0.07), and there were also no significant differences in partner preference between time periods of PPT (48 and 72 hours) (*U* = 119, *p* = 0.75, *effect size r* = 0.06) (see Supplementary Data 1).

### Male and female voles share network-level changes related to social bonding

Network Based Statistics analysis yielded significant changes between baseline, 24h and 2 weeks of cohabitation MR sessions (*p_FWE_*= 0.011) in a network consisting of nine regions: ACC, AON, dHIP, LS, mPFC, PVN, RSC, vHIP, and VTA (Figure 3a). No significant differences were found between male and female voles. The nodes with more changes detected in their connections were the LS, with four, and the ACC with three. Post-hoc analyses identified differential longitudinal changes among rsfMRI sessions. Specifically, eight connections had significant changes after 24 hours of cohabitation: ACC-vHIP (*p_FDR_*= 0.0014) and LS-RSC (*p_FDR_*= 0.0049) showed increased functional connectivity, while the ACC-VTA (*p_FDR_*= 0.0007) and the PVN-RSC (*p_FDR_*= 0.0452) had decreased connectivity in the same period of time. Both ACC-vHIP and ACC-VTA had no differences between session 3 (after 2 weeks of cohabitation) and baseline (before cohabitation), suggesting acute plastic changes related to 24 hours of cohabitation; however, LS-RSC (*p_FDR_*= 0.0113) and LS-mPFC (*p_FDR_*= 0.0094) have increased connectivity, while ACC-LS (*p_FDR_*= 0.0046) has decreased connectivity between sessions1 and 3. In addition, five connections reflected differences between 24h and 2 weeks of cohabitation (sessions 2 and 3): ACC-LS (*p_FDR_*= 0.0046), ACC-mPFC (*p_FDR_*= 0.0673), and AON-LS (*p_FDR_*= 0.0190) had decreased connectivity, but LS-mPFC (*p_FDR_*= 0.0003) and mPFC-dHIP (*p_FDR_*= 0.0015) exhibit increased connectivity in session 3, suggesting long-term functional changes related to cohabitation (Figure 3, b-i).

**Figure 3.**
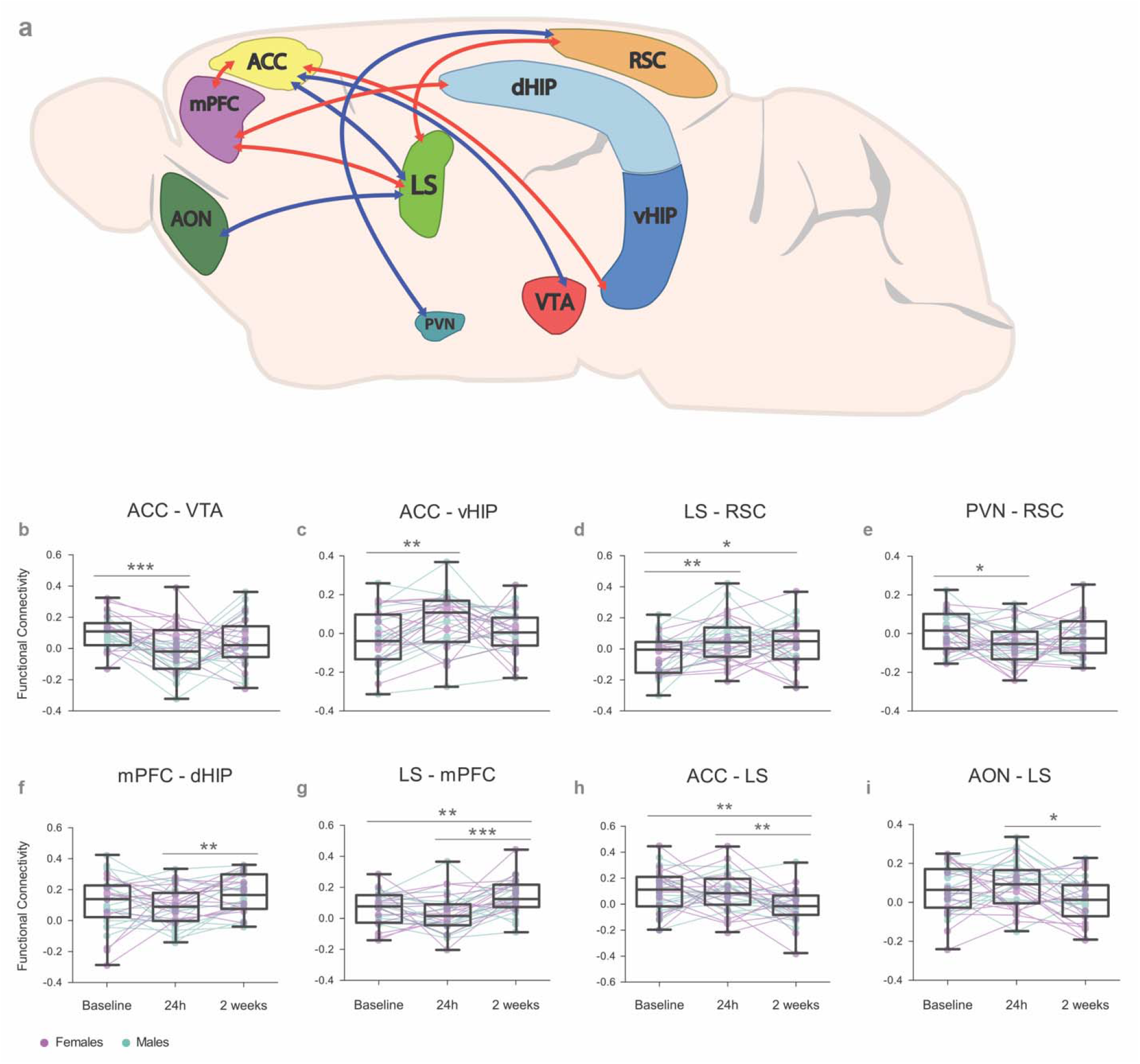
Changes in brain functional connectivity before and after cohabitation with mating in female and male prairie voles: **a)** NBS analysis results represented in a prairie vole brain with regions (nodes) comprising the brain network that undergoes significant changes in functional connectivity after cohabitation with mating. Interregional connectivity (edges) is shown by color code. Red: increase of functional connectivity; blue: decrease of functional connectivity. ACC: anterior cingulate cortex. AON: anterior olfactory nucleus. dHIP: dorsal hippocampus. LS: lateral septum. mPFC: medial prefrontal cortex. PVN: paraventricular nucleus of the hypothalamus. RSC: retrosplenial cortex. vHIP: ventral hippocampus. VTA: ventral tegmental area. **(b-i)** Functional connectivity values displayed in boxplot graphs (25 and 75 percentile) with whiskers (min and max) data points. Connecting lines track longitudinal data of each subject between regions through specific time-points (MR sessions): Baseline, 24h and 2 weeks of cohabitation. Color codes of connecting lines distinguish male (green) from female subjects (violet). False discovery rate (FDR) post-hoc significant differences are shown: *<.05, **<.01 ***<.001.

### Baseline functional connectivity predicts the display of affiliative behavior

Since behavioral analysis during cohabitation reflected a high level of individual variation, we tested if functional connectivity had a relationship in the display of huddling, an affiliative behavior shown to be socially relevant in the prairie vole and that may influence pair bond induction and formation (Amadei et al., 2017). Hence, we assessed the relationship between baseline functional connectivity, i.e. before cohabitation, and huddling latencies during cohabitation in voles of both sexes (*N*=30). NBS analysis found significant negative linear relations in the following sub-networks: ACC-NAcc-BLA-DG (*p* = 0.013), RSC-VTA (*p* = 0.038), and MeA-VP (*p* = 0.041) (Figure 4). Our results show that the higher the connectivity between these regions before cohabitation, the shorter the huddling latencies during cohabitation in voles of both sexes. Additionally, *a posteriori* Pearson correlations confirmed the correlation strength between each connected node in the ACC-NAcc-BLA-DG sub-network: ACC-NAcc (r(28) = −0.493, *p* = 0.0057), NAcc-BLA (r(28) = −0.559, *p* = 0.0013), and BLA-DG (r(28) = −0.425, *p* = 0.0193) (Figure 4, a-e). These results show that functional connectivity between these regions reflect the predisposition of voles to display affiliative behavior, in both females and males.

**Figure 4.**
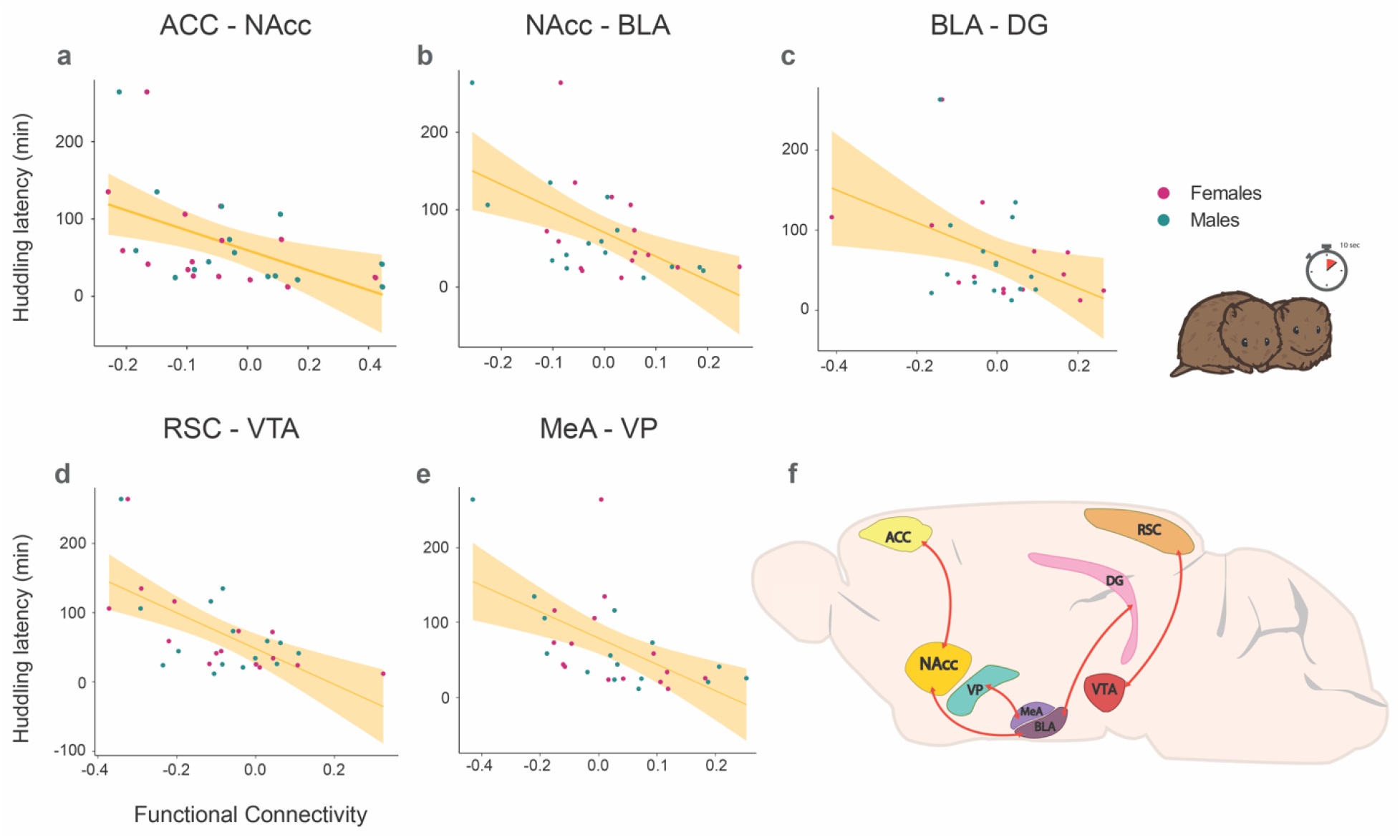
Relationships between baseline functional connectivity and affiliative behavior (huddling) during cohabitation with mating in male and female prairie voles. Scatterplot graphs **(a-e)** showing significant interregional negative correlations (with best line fit) between baseline functional connectivity and huddling latencies (minutes) during cohabitation. The higher the connectivity between these regions before cohabitation, the shorter the huddling latencies during cohabitation in voles of both sexes. **(f)** Representation of a prairie vole brain with regions (nodes) comprising brain sub-networks that correlate with huddling latency. ACC: anterior cingulate cortex. BLA: basolateral amygdala. DG: dentate gyrus. MeA: medial amygdala. NACC: nucleus *accumbens*. RSC: retrosplenial cortex. VP: ventral pallidum. VTA: ventral tegmental area.

### Partner preference measured between 48h and 72h of cohabitation predicts long-term functional connectivity between MeA and VP

Given that several network-level changes were found during cohabitation, we evaluated if functional connectivity could be related to the level of partner preference measured at 48h and 72h of cohabitation. NBS analysis found a significant positive linear relationship between the partner preference index and the MeA-VP connectivity at two weeks of cohabitation (*p*=0.038, *N*=30) (Figure 5). This result suggests that the higher the partner preference is between 48h and 72h of cohabitation, the higher functional connectivity will be between MeA and VP after two weeks of cohabitation.

**Figure 5.**
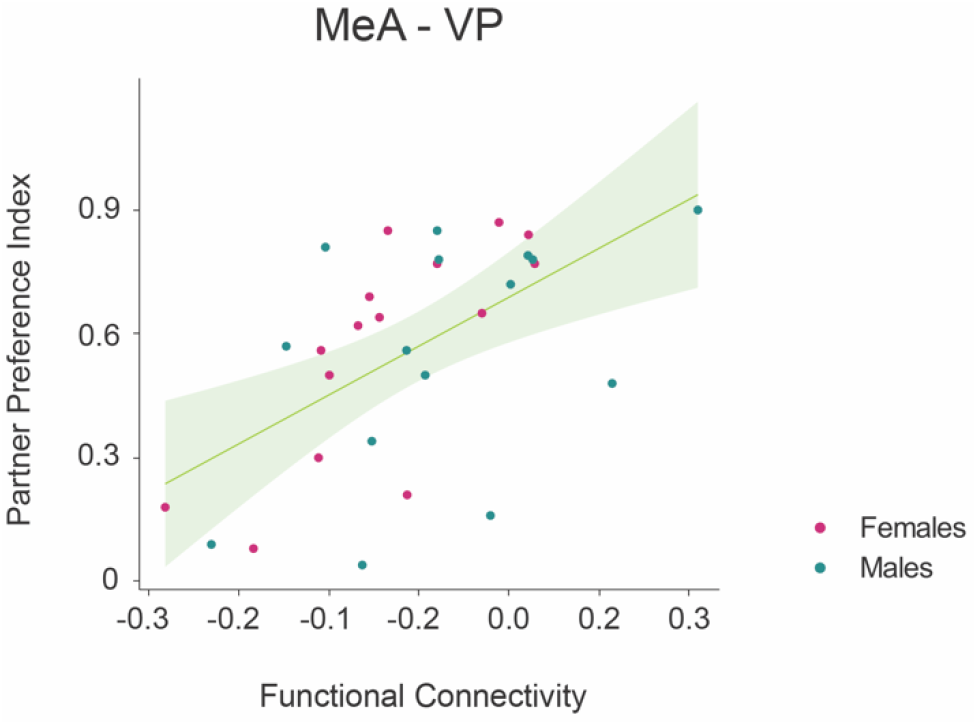
The level of partner preference at 48 and 72h of cohabitation predicts long-term functional connectivity between the medial amygdala and the ventral pallidum. Scatterplot graph with best line fit of partner preference index and functional connectivity at 2 weeks of cohabitation, suggesting that partner preference predicts levels of functional connectivity at 2 weeks of cohabitation.

## Discussion

Several studies have described the relevance of different brain regions involved in pair bond induction and maintenance in prairie voles (Johnson & Young, 2017; Walum & Young, 2018). However, longitudinal explorations of the brain before and after pair bonding are scarce (Bales et al., 2007), especially from a network perspective. Here, by using rsfMRI, we were able to detect a brain network with significant longitudinal changes in the functional connectivity of prairie voles after pair bonding. Post-hoc analyses revealed differential short- and long-term connectivity changes mainly involving ACC, LS and the hippocampus. Furthermore, baseline functional connectivity (before cohabitation) predicted the latency for huddling during the first hours of cohabitation, providing the potential neurofunctional substrate for the variability in affiliative behavior and further extending the recent findings that the corticostriatal electric activity modulates social bonding in prairie voles (Amadei et al., 2017). Finally, partner preference, a potential measure of pair bonding evaluated at 48 or 72 hours of cohabitation, predicted long-term functional connectivity between the medial amygdala and the ventral pallidum. The connectivity between the latter regions at baseline also predicted affiliative behavior, further highlighting the relevance of such interaction for both initial affiliative behavior and long-term pair bonding. We further discuss the potential neurophysiological basis and implications of our findings.

### Longitudinal changes in functional connectivity after pair bonding

It has been proposed that pair bonding results from the convergence of the mesolimbic dopamine reward circuit and social discrimination circuits (Walum & Young, 2018). The results here presented are consistent with this model, by demonstrating both short and long-term changes in a brain network including regions largely associated with reward/motivation (ACC, VTA, NAcc) and the social decision making network, specifically related to sensory contextualization (LS, RSC, dHIP), saliency processing (LS, ACC, VTA), memory formation and retrieval (vHIP, dHIP, mPFC) and in neuronal regions involved in OXT synthesis such as the PVN.

Changes in functional connectivity after 24 hours of cohabitation may result from familiarization of a new spatial context that also implies a novel social context, i.e. exposure to new housing and to a novel, opposite-sex conspecific stranger. During this process, subjects are exposed to novel stimuli and engage in their first socio-sexual interactions, forming new memories of the partner. Furthermore, functional connectivity changes detected at 2 weeks of cohabitation may be related to long-term modulation of behavior as a result of social bonding, in which the partner and its associated cues become salient/rewarding, and after-bonding behaviors such as mate and territorial guarding may appear. In general, these temporally-dynamic functional connectivity changes may be related to the modulation of socio-sexual interactions with the partner. While the nodes here identified are interconnected into a larger network and their precise contribution to complex social behavior remains elusive to our methods, their potential role will be discussed based on previous results and the connectivity patterns here identified.

The functional interactions that had changes at 24 hours of cohabitation included the ACC and VTA, which may regulate saliency of stimuli. The ACC has been considered an important region in the decision-making process between sensory perception, motivation and final motor performance (Assadi et al., 2009). On the other hand, the VTA is a region known to modulate motivational, reward, and emotionally salient stimuli (Gunaydin et al., 2014). There is evidence that the VTA modulates DA release in prefrontal regions to regulate sexually relevant stimuli (Cortes et al., 2019). Therefore, it is likely that this functional connectivity change may be related to a regulation in sexual motivation as a result of recently-acquired sexual experience. This initial motivation may be necessary to promote further socio-sexual interaction and the likely induction of pair bonding.

Social interaction would necessarily require new memory consolidation, and possibly, formation of the neural representation of the partner. Our data showed functional connectivity changes occurred in the hippocampus after 24 hours of cohabitation (ACC-vHIP, Figure 3), which has demonstrated to be critical in encoding spatial and mnemonic information in rodents (Lin et al., 2017). ACC-vHIP activity has been previously associated with the regulation of fear in novel environments and in contextual fear generalization in rodents (Bian et al., 2019). However, the ventral hippocampus has also shown to contribute in social memory processing (Chiang et al., 2018). Additionally, the ACC is thought to facilitate the integration of new information into existing internal representations that can motivate and modify future behavior (Kolling et al., 2016). Thus, increased ACC-vHIP connectivity after 24 hours of cohabitation might be related to social memory formation, but it may also be related to potentially increased anxiety triggered by a new spatial and social environment.

At 24 hours of cohabitation, an increase of connectivity between the LS and the RSC was also identified. The LS is a structure involved in social recognition and retrieval of relevant social information (Bielsky et al., 2005). In regard to the RSC, it is known to be involved in the processing of spatial memory and cognition in rodents (Todd & Bucci, 2015). The increased connectivity between the LS and the RSC at 24 hours may be related to the processing of contextual information and in associating social information with the spatial context. A significant decrease of connectivity was also detected between the RSC, which is sensitive to OXT, and the PVN, a region known for OXT synthesis and release (Jurek & Neumann, 2017).

After 2 weeks of cohabitation, functional connectivity between the mPFC and the dHIP was increased. The dorsal region of CA2 has been characterized as a hub for sociocognitive memory processing in rodents, specifically for social memory encoding, consolidation, recall (Meira et al., 2018), and social recognition (Hitti & Siegelbaum, 2014). Also, the mPFC has been proposed as a critical region for remote memory retrieval and for memory consolidation, reportedly relying on the hippocampus (Euston et al., 2012). Thus, their increased interaction could support the integration of new memories into pre-existing networks (Preston & Eichenbaum, 2013). Potentially, the increased connectivity between these regions may be related to the incorporation of the partner to long-term memory representations. Indeed, OXT receptor activation in dHIP promotes the persistence of long-term social recognition memory in mice (Lin et al., 2017).

A long-term increase of connectivity was detected between the LS with the mPFC and the RSC, and decreases of connectivity were observed between ACC-LS and AON-LS at 2 weeks of cohabitation. As mentioned before, the LS is involved in social recognition and retrieval of relevant social information. In rats, it has been shown that OXT release in the LS is required for the maintenance of social memory, modulated by the relevance of the social stimulus (Lukas et al., 2013). Thus, LS connectivity changes might be related to long-term social recognition and behavior modulation, mediated by afferents from prefrontal structures, i.e. the mPFC and the ACC (ACC-LS-mPFC), and by sensory-related information from the AON (AON-LS) and contextual information (LS-RSC). Indeed, the aforementioned ACC may have a role in saliency regulation and in decision-making processes, while the mPFC is proposed to have a role in the initiation, maintenance and modulation of social attachment and behavior in rodents (Ko, 2017; Tanimizu et al., 2017). On the other hand, AON activation is essential for conspecific recognition and promotes the extraction of relevant olfactory information from the MOB in rats (Oettl et al., 2016), and as previously noted, the RSC may contribute with the relationship between spatial information with a social environment (Todd & Bucci, 2015). Since AVP and OXT receptor blockade in the LS inhibits, while AVP administration enhances pair bond formation in prairie voles (Liu et al., 2001), and male voles exposed to females have increased Fos-immunoreactive cells in the LS (Wang et al., 1997); it is enticing to propose that the LS is the node that enables the integration of olfactory information, with mnemonic and motivational/saliency information, which are necessary for conspecific recognition and subsequent modulation of behavior.

Although both AVP and OXT have been proposed to have sex-differentiated roles in pair-bond formation (L. J. Young & Wang, 2004), the functional connectivity here explored showed no significant differences between sexes, nor significant sex-session interactions. Accordingly, our results have shown modulation of consistent functional networks for both female and male prairie voles. We propose that the longterm functional connectivity changes observed in the network are related to social bonding and may lead to pair bonding induction and maintenance in both female and male prairie voles.

### Correlations between sub-network functional connectivity with huddling latencies and Partner Preference Index

Even though it’s been reported that in prairie voles 24 hours of cohabitation or 6 hours of *ad libitum* mating are sufficient for a pair bond to be developed (Williams et al., 1992), a considerable amount of evidence has shown that other factors influence its development and maintenance. Specifically, AVP (Ophir et al., 2008) and OXT receptor gene expression and density (King et al., 2016), paternal nurturing (Ahern & Young, 2009) or deprivation (Barrett et al., 2015), have shown to produce variability in the exhibition of prairie vole social behavior. Subjects of both sexes in this study showed a wide variability in their bonding behavior evaluated between 48 and 72h of cohabitation, even though they were under the same experimental conditions. It is likely that the sum of previously mentioned factors gives each subject a distinctive brain network configuration that ultimately relates to bonding behavior. Thence, we hypothesized there may be individual differences in functional connectivity that could explain the variability in socio-sexual behavior. Indeed, we identified three functional networks for which the functional connectivity at baseline was negatively related to huddling latencies during the first hours of cohabitation. In other words, baseline functional connectivity predicts how quickly subjects would begin affiliative huddling with an opposite sex conspecific. Huddling is a measurable affiliative behavior in prairie voles and a useful indicator of social receptiveness (Salo et al., 1993). The larger network included four nodes, related as follows: ACC-NAcc-BLA-DG (Figure 4), while the other two only included two nodes: MeA-VP and VTA-RSC. These subnetworks involve regions reported to have a role in social memory and recognition, spatial, and reward-seeking mechanisms. Specifically, the BLA and the ACC have been found relevant in coordinating brain activity when social interaction is initiated and in the formation of social recognition memory through gene expression (Tanimizu et al., 2017). In line with our results, the BLA may act as an associative site for stimulus-outcome representations (Cardinal et al., 2002), allowing an appropriate response according to previous social encounters, and may require mnemonic information encoded by the DG (BLA-DG). Part of a social interaction, once a conspecific has been recognized, involves a reward component. The NAcc is known to translate reward-predictive information from the amygdala (NAcc-BLA) to promote cue-evoked, reward-seeking behavioral responses in rodents (Ambroggi et al., 2008); additional input from the ACC (ACC-NAcc) would be necessary for a social decisionmaking process and motor performance (Assadi et al., 2009). Furthermore, a recent study showed that in female prairie voles, the functional connectivity of the prefrontal cortex and NAcc after the first encounter predicts affiliative huddling towards a partner, and the activation of such circuit biases later preference towards a partner (Amadei et al., 2017). Our results extend such findings, showing that a larger network including similar corticostriatal connectivity, measured even before the exposure to a potential partner, predicts affiliative behavior. Moreover, such relation is consistent for both male and females, and the circuit includes the amygdala and hippocampus, as predicted by Amadei and cols. (2017).

In addition, two other circuits also predicted future affiliative behavior, the VTA-RSC and MeA-VP (Figure 4). The former may indicate a potential bias for a more efficient integration of socially-rewarding stimuli with contextual information of the partner (Hung et al., 2017; Todd & Bucci, 2015). While in the latter, it is known the VP plays a major role in reward and motivation (Smith et al., 2008), encoding reward value earlier and more robustly than the NAcc and potentially being a crucial downstream mediator of NAcc reward-related functions (Ottenheimer et al., 2018), including pair bonding (M. M. Lim & Young, 2004). Since MeA activity in rodents is necessary for social recognition (Ferguson et al., 2001) and responds to sex-specific chemosensory cues (Wang et al., 1997; Yao et al., 2017), increased MeA-VP connectivity may improve the rewarding response of social and chemosensory cues from the partner.

Moreover, the MeA-VP connectivity after 2 weeks of cohabitation was also predicted by the Partner Preference Index at 48 and 72 hours, which captures the strength of a pair bond (Williams et al., 1992). This relation is not surprising given the recent evidence that the MeA synchronizes socio-sexual behavior by the enhancement of preference for a sexual partner (Adekunbi et al., 2018). Meanwhile, the VP is related to the hedonic or motivational impact of the partner-related stimuli (Smith et al., 2008). The identified relation suggests the strengthening of this sub-network (MeA-VP) along 2 weeks of cohabitation in voles with higher partner preference between 48 and 72 hours of cohabitation (Figure 5).

We found particularly interesting that the NAcc, MeA, VP and BLA, although deemed relevant for social bonding, are not nodes of the main network with longitudinal changes across rsfMRI sessions. However, the connectivity of these regions actually relates to the affiliative behavior of the prairie voles, highlighting the role of their neural circuits in the variability of prairie vole social behavior, i.e. “wanderer” or “resident” mating tactics; in which resident voles maximize mating success through mate guarding and reduced territory space, in contrast to wanderers that overlap many home ranges to increase opportunistic mating (Getz et al., 2005). This is supported by previous work demonstrating that AVP receptor expression in the RSC predicts sexual fidelity and territorial behavior in male prairie voles (Ophir et al., 2008). In addition, AVP receptor overexpression in VP induced partner preference in socially promiscuous meadow voles (Lim, Wang, et al., 2004), and variation in OXT receptor density in the NAcc relates to partner preference in prairie voles (King et al., 2016). Consequently, sub-network functional connectivity variability may also reflect behavioral diversity influenced by genetic and environmental factors.

The prairie vole has proven to be a valuable model that has enabled the characterization of the neurobiological mechanisms behind complex socio-sexual behaviors, potentially useful to understand human social bonding and its alterations in psychological disorders. To our knowledge, this is the first study to demonstrate both short and long-term changes in the functional connectivity of multiple interacting brain regions (networks) of prairie voles in cohabitation with mating. However, there are some limitations related to the use of rsfMRI. First, the animals were scanned under anesthesia, which may potentially alter the brain functional connectivity. Yet, we have used an anesthesia protocol that practically preserves the functional interactions as in the awake state (Grandjean et al., 2014, Paasonen et al., 2018). Although functional neuroimaging of awake prairie voles is possible, the acclimation process may induce significant stress and even increase the risk of physical harm (Yee et al., 2016), limiting the longitudinal design and the interpretation of the results. Second, the precision in the definition of the ROIs is limited by the shape and the size of the voxels, being hard to assure that the defined regions specifically and uniquely include signal from the anatomical regions of interest. Nevertheless, we are certain that the defined ROIs include the actual regions of interest, and at the least, the changes here reported are related to those areas and their surrounding tissue. It is important to note that functionally connected nodes may not necessarily have direct axonal projections to each other, and the interaction between nodes could be mediated or relayed through other structures (Friston et al., 1993). Also, contrary to other electrophysiological or neuropharmacological methods, functional MRI captures an indirect measure of neuronal activity (Kim & Ogawa, 2012). However, it poses the great advantage that it allows the longitudinal exploration of the brain, with the best spatial resolution and wider coverage that any imaging method can achieve non-invasively. The consistency of our results with recent findings using direct electrophysiology readings (Amadei et al., 2017) further support their relevance in identifying the neurophysiology of complex social behaviors.

## Conclusions

Our findings suggest there is a brain network that encompasses short- and long-term changes in the socio-sexual behavior of prairie voles as a consequence of social bonding and mating during cohabitation. Such functional connectivity changes may be involved in the process of pair bonding and in the formation and consolidation of the neuronal representation of the partner. Furthermore, the functional connectivity of specific subnetworks including the ACC, NAcc, BLA, DG, MeA, VP, RSC and VTA, predict huddling latencies, suggesting there is a neurobiological predisposition to social bonding, while another network (MeA-VP) correlates with the level of early partner preference. In summary, our findings suggest: 1) baseline functional connectivity predicts the level of affiliative behavior even before sexual experience and, 2) sexual experience and long-term cohabitation induces network-wide changes in socio-sexual relevant circuits. Functional connectivity may also aid in exploring the mechanisms that underlie individual variability in the expression of socio-sexual behavior. Overall, our findings have shown network-level changes associated with the process of social bonding and provide a novel approach to further investigate the neurophysiology of complex social behaviors displayed in the prairie vole.

## Acknowledgements

We thank Dr. Fernando A. Barrios for his valuable comments to the manuscript, and Deisy Gasca, Martín García, Alejandra Castilla, Leopoldo González Santos, Nuri Aranda López and Ma. De Lourdes Lara for their technical assistance. The authors thankfully acknowledge the imaging resources and support provided by the Laboratorio Nacional de Imagenología por Resonancia Magnética (LANIREM), part of the CONACYT’s network of national laboratories. This research was supported by grants CONACYT 252756 to WP, 253631 to RGP, UNAM-DGAPA-PAPIIT IN202818 to WP, IN212219-3 to SA, IN203518-3 to RGP, INPER 2018-1-163 to NFD. LJY’s contribution was supported by NIH grants P50MH100023 to LJY and P51OD011132 to YNPRC. M.F. López-Gutiérrez was supported by a fellowship from CONACYT (fellowship #626152) for her Masters of Science studies in Neurobiology at UNAM (Maestría en Ciencias [Neurobiología], UNAM).

